# Gene co-expression patterns in Atlantic salmon adipose tissue provide a molecular link among seasonal changes, energy balance and age at maturity

**DOI:** 10.1101/2023.08.30.555548

**Authors:** Ehsan Pashay Ahi, Jukka-Pekka Verta, Johanna Kurko, Annukka Ruokolainen, Pooja Singh, Paul Vincent Debes, Jaakko Erkinaro, Craig R. Primmer

## Abstract

Sexual maturation in many fishes requires a major physiological change that involves a rapid transition between energy storage and usage. In Atlantic salmon, this transition for the initiation of maturation is tightly controlled by seasonality and requires a high-energy status. Lipid metabolism is at the heart of this transition since lipids are the main energy storing molecules. The balance between lipogenesis (lipid accumulation) and lipolysis (lipid use) determines energy status transitions. A genomic region containing a transcription co-factor of the Hippo pathway, *vgll3*, is the main determinant of maturation timing in Atlantic salmon. Interestingly, *vgll3* acts as an inhibitor of adipogenesis in mice and its genotypes are potentially associated with seasonal heterochrony in lipid storage and usage in juvenile Atlantic salmon. Here, we explored changes in expression of more than 300 genes directly involved in the processes of adipogenesis, lipogenesis and lipolysis, as well as the Hippo pathway in the adipose tissue of immature and mature Atlantic salmon with distinct *vgll3* genotypes. We found molecular evidence consistent with a scenario in which immature males exhibiting *vgll3* genotype specific seasonal direction of change in their lipid profiles. We also identified components of the Hippo signaling pathway as potential major drivers of *vgll3* genotype specific differences in adipose tissue gene expression. This study demonstrates the importance of adipose gene expression patterns for directly linking environmental changes with energy balance and age at maturity through genetic factors bridging lipid metabolism, seasonality and sexual maturation.

## Introduction

Ecological factors play fundamental roles in the maintenance of variation in life-history genes (Jops & O’Dwyer, 2023), particularly those linked with “pace-of-life” traits (Arnqvist & Rowe, 2023). The timing of sexual maturation (age at maturity) is a crucial life-history trait influenced by environmental cues as well as genetic and physiological mechanisms regulating the onset of puberty (Taranger et al., 2010). Age at maturity also subsequently affects other crucial fitness traits such as survival and reproductive success (Mobley et al., 2021). Elucidating the molecular basis of variation in age at maturity, especially in relation to environmental cues, is thus a pivotal step towards understanding the maintenance of the diversity in life-history traits and its ecological significance.

Sexual maturation in Atlantic salmon is tightly linked with seasonal environmental changes. For example, temperature changes can affect metabolism and body condition, via lipid allocation, by impacting food availability, and thereby growth rate (House, Debes, Kurko, Erkinaro, & Primmer, 2023). Lipid allocation has been found to be especially important for age at maturity in Atlantic salmon due to their need to reach a certain threshold of lipid reserves in order to initiate maturation (Rowe, Thorpe, & Shanks, 2011). This allocation process can be strongly affected by variation in replenishing reserves across seasonal changes, including periods of low temperatures and food availability (Mogensen & Post, 2012). In Atlantic salmon, a single locus including the gene vestigial-like family member 3 (*vgll3*) is the major genetic determinant of maturation timing (explaining over 39% of the variation in the age at maturity) (Barson et al., 2015; Czorlich, Aykanat, Erkinaro, Orell, & Primmer, 2018) and the effects of *vgll3* on age at maturity can be observed in males as early as one year of age in controlled conditions (Debes et al., 2021; Sinclair-Waters et al., 2021; Verta et al., 2020). Interestingly, the human ortholog (*VGLL3*) has also been shown to be associated with age at maturity (Cousminer, Widén, & Palmert, 2016; Perry et al., 2014). In addition to being linked with maturation age, *vgll3* has also been shown to function as an inhibitor of adipogenesis in mice (Halperin, Pan, Lusis, & Tontonoz, 2013). In salmon, energy storage in individuals with the distinct *vgll3* genotypes varies across seasons thereby providing an interesting link between energy expenditure and environmental change (House, Debes, Kurko, et al., 2023).

Recent studies of one year old Atlantic salmon identified strong association between *vgll3* early (E) and late (L) maturation alleles and the expression of key reproductive axis genes (Ahi, Sinclair-Waters, Moustakas-Verho, Jansouz, & Primmer, 2022). Gene co-expression network analysis predicted that the *vgll3* genotype effects on reproductive axis genes are likely to be mediated by differential activation of the Hippo signaling pathway (Ahi, Sinclair-Waters, Donner, & Primmer, 2023). The Hippo pathway is emerging as a key molecular signal not only in controlling sexual maturation in vertebrates (Kjærner-Semb et al., 2018; Kurko et al., 2020; Sen Sharma, Vats, & Majumdar, 2019), but also in balancing adipocyte proliferation versus differentiation, and the activity of Yap1 (a major transcription co-factor of the Hippo pathway) has been found to be indispensable during adipogenesis (Ardestani, Lupse, & Maedler, 2018). Strikingly, Hippo pathway appears to be an essential molecular cue for responding to environmental effects, such as changes in diet fat and temperature, at transcriptional levels (Luo et al., 2020; Shu et al., 2019). Vgll3 acts as a major activating transcription co-factor of the Hippo pathway and competes with the other major transcription co-factor of the pathway, Yap1, which mainly acts as an inhibitor of the Hippo pathway (Hori et al., 2020; Kurko et al., 2020). Together, this suggests that Atlantic salmon, with distinct *vgll3* alleles tightly linked to maturation and adipogenesis processes, are an excellent natural model system to explore molecular mechanisms directly linking sexual maturation, energy acquisition and environmental changes.

In this study, we used a custom-made Nanostring gene expression panel to investigate the expression patterns of 337 genes including genes encoding adipogenesis related factors, Hippo pathway components and their associated interacting partners/signals in the adipose tissue of mature and immature male Atlantic salmon with *early* (E) or *late* (L) maturing *vgll3* genotypes.

Comparisons between maturity status and *vgll3* genotype enabled application of pathway– and regulatory network-based approaches for better understanding underlying molecular processes.

## Materials and methods

### Fish material and tissue sampling

Individuals used in the study were from the same population (Oulujoki) and cohort used in Verta et al. (2020). This provided access to individually passive integrated transponder tagged individuals with known *vgll3* genotypes (see Verta et al., (2020) for more details of crossing and rearing). For this study, male individuals were sampled at the ages of 1.5 to 2 years post fertilization (Debes, Piavchenko, Erkinaro, & Primmer, 2020). Following euthanization with an overdose of MS222, various tissues, including visceral adipose tissue, of individual males were sampled at three time points, each reflecting a different maturation development stage:

*Immature 1*-individuals were sampled in late spring (5-21 May), average mass 17.5 g (range 10.2-27.4 g), average length 12.1 cm (range 9.1-13.6 cm), and showed no indications of gonad development (gonadosomatic index; GSI = 0).

*Immature 2*-individuals were sampled in the summer (4-17 July), average mass 33.7 g (range 21.1-76.6 g), average length 17.3 cm (range 14-18.5 cm), and some individuals showed initial signs of the commencement of phenotypic maturation processes (GSI = 0.0-0.4).

*Mature*-individuals were sampled during the expected spawning period in the early autumn (1-15 October), average mass 82.7 g (range 60.3-120.6 gm), average length 20.8 cm (range 16.4-22.7 cm), and all individuals had well developed gonads reflective of being mature (GSI > 3).

These tiss es were lash ro en in li id nitro en and stored at – 0 until processing.

### RNA extraction

In total, RNA was extracted from 32 samples of male visceral adipose tissue using a NucleoSpin RNA kit (Macherey-Nagel GmbH & Co. KG). The samples were transferred to tubes with 1.4 mm ceramic beads (Omni International), Buffer RA1 and DDT (350ul RA1 and 3,5ul DDT 1M) and homogenized for 2 min (6 x 20 s) at 30 Hz using Bead Ruptor Elite (Omni International). The RNA extraction steps were conducted accordin to the man act rer’s instr ctions. The kit also contained a built-in DNase step to remove residual gDNA. At the end, RNA extracted from each sample was eluted in 50 µl of nuclease free water. RNA quantity was measured with NanoDrop ND-1000 (Thermo Scientific, Wilmington, DE, USA) and quality was assessed with 2100 BioAnalyzer system (Agilent Technologies, Santa Clara, CA, USA), and the RNA integrity number (RIN) was above 7 in all the samples. From each extraction, 100 ng of total RNA was used for hybridization step in the Nanostring panel.

### Nanostring nCounter mRNA expression panel

NanoString nCounter is a multiplex nucleic acid hybridization technology that enables reliable and reproducible assessment of the RNA expression of up to several hundred genes in a single assay (Goytain & Ng, 2020). This technology has several advantages for ecological and evolutionary research as it requires very small amount of RNA input with lower quality than RNA-seq, does not have an amplification step and it can detect very low RNA expression levels. The Nanostring panel of probes used in the current study was an extended panel from that used in (Kurko et al., (2020) that was previously developed to investigate age at maturity related gene expression in Atlantic salmon (more than 100 genes added). These genes include an extensive list of Hippo pathway components and other known genes having direct cross-talk with the Hippo pathway. The selection was conducted based on the literature and the IPA (Ingenuity Pathway Analysis) tool (Qiagen) and other freely available web tools and databases (Kurko et al., 2020). In addition, the panel contains probes for the age-at-maturity-associated genes in Atlantic salmon; *vgll3a* on chromosome 25 and *six6a* on chromosome 9, and their corresponding paralogs *vgll3b* on chromosome 21 and *six6b* on chromosome 1. Further, probes for other functionally associated genes with important roles in metabolism (e.g. lipidogenesis), cell fate (adipogenesis) and sexual maturation (e.g. HPG axis), were also included. Most of the candidate genes possess one or more paralogs due to the a recent Atlantic salmon genome duplication, and therefore, paralogs of each gene of interest were also included after identification through the SalmoBase (http://salmobase.org/), and NCBI RefSeq databases. Further details on the selection of the genes/paralogs and naming of them can be found in Kurko et al., 2020. Gene accession numbers, symbols, full names and functional categories are listed in Supplementary Table. The mRNA expression levels of the candidate genes were investigated using Nanostring nCounter Analysis technology (NanoString Technologies, Seattle, WA, USA). Probes for each gene paralog, targeted at all known transcript variants, were designed using reference sequences in the NCBI RefSeq database. However, for some genes, it was not possible to design paralog-specific probes, as sequence similarity between paralogs was too high. The RNA samples were analyzed using nCounter Custom CodeSet for probes and nCounter Master kit (NanoString Technologies). The RNA of each sample was denatured, hybridized with the probes overnight and in the following day post-hybridization purification and image scanning were conducted.

### Data analysis

Among nine candidate reference genes in the panel, seven genes, including *ef1a* paralogs (*ef1aa*, *ef1ab* and *ef1ac*), *hprt1*, *prabc2* paralogs (*prabc2a* and *prabc2b*) and *rps20*, were selected for data normalization since they showed a low coefficient of variation (CV) values across the samples. The excluded reference genes were *actb* and *gapdh* showing very high variation (CV% > 100), and even though they are commonly used as reference genes in many studies, they seemed to be unsuitable for data normalization in the adipose tissue of Atlantic salmon. Subsequently, the raw count data from the Nanostring nCounter mRNA expression was normalized by RNA content normalization factor for each sample calculated from geometric mean count values of the selected seven reference genes. After normalization, a quality control step was conducted on the data and all the samples passed the default limit based on the nSolver Analysis Software v4.0 (NanoString Technologies; www.nanostring.com/products/nSolver). Mean of the negative controls were subtracted while analyzing the data using the software and positive control normalization was performed using the geometric mean of all positive controls as recommended by the manufacturer. A normalized count value of 20 was set as a background signal threshold, and consequently, 121 genes had on average below background signal across the samples, which left 202 genes to be considered for further analyses. The differential expression analysis was performed using the log-linear and negative binomial model (lm.nb function), as implemented in Nanostring’s nSolver Advanced Analysis Module (nS/AAM). The maturation status and genotypes were selected as predictor covariates in the model as suggested by nS/AAM. Multiple hypothesis adjustment was performed using the Benjamini-Yekutieli method (Benjamini & Yekutieli, 2001) within the software and adjusted p-values < 0.05 were considered significant (Supplementary table). The log-transformed expression values were also used for calculations of pairwise Pearson correlation coefficients (r) between the expression of each gene and GSI values across all the samples.

The Weighted Gene Coexpression Network Analysis (WGCNA version 1.68) R-package (version 5.2.1) was implemented to identify gene co-expression networks (GCN) (Langfelder & Horvath, 2008). Since our main interest was in the comparison of alternative *vgll3* genotypes, all samples from both maturation statuses and time-points within each genotype were used as biological replicates, providing sufficient statistical power for WGCNA. To identify sample relationships, we conducted hierarchical clustering of samples based on gene expression. Coexpression networks were constructed with the following steps: (1) calculation of Pearson correlation coefficients to identify and measure gene coexpressions, (2) calculation of an adjacency matrix (with reference to a scale free topology) using the coefficients (3) calculation of the topological overlap distance matrix using the adjacency matrix, (4) hierarchically cl sterin o enes (method = avera e) sin the topological overlap distance, (5) identification of coexpressed modules of genes using the cutTreeDynamic function with a minimum module size of 10 genes (6) color assignment for each module and representation module-specific expression profile by the first principal component (module eigengene) of each module, and (6) merging highly similar modules based on module eigengene (ME) dissimilarity (distance threshold of 0.25) in order to find the final set of coexpressed gene modules. Next, a conditional coexpression analysis was set (as described by Singh et al., 2021); i.e. coexpression networks were constructed for each *vgll3* genotype separately, to identify the preservation of EE modules in the LL network and vice versa. A softpower of 7 was used to construct adjacency matrix. Finally the module preservation statistics were calculated using WGCNA to check how the density and connectivity of modules defined in the reference dataset (e.g. EE) were preserved in the query dataset (e.g. LL) (Langfelder, Luo, Oldham, & Horvath, 2011). A permutation test was implemented to repeatedly permute genes in the query network in order to calculate Zscore. Individual Z scores from all permutations (200) were summarised as a Zsummary statistic.

Gene ontology (GO) enrichment analysis was conducted to identify overrepresented biological process in each GCN using Manteia (Tassy & Pourquié, 2014). The GO enrichment criteria were limited to FDR < 0.05 with GO speci icity level o 2 set as a cut-off. To predict potential gene interactions and key genes with the highest number of interactions (interacting hubs), the identified differentially expressed genes in each comparison were converted to their conserved orthologs in human (which have the highest amount of validated/studied interactome data across vertebrates) and used as input for STRING version 12.0, one of the largest freely available knowledge-based interactome database for vertebrates (Szklarczyk et al., 2023). The interaction predictions between genes were based on data from structural similarities, cellular co-localization, biochemical interactions and co-regulation. The confidence level to predict each interaction/molecular connection was set to medium (the default).

## Results

### Differences in gene expression between *vgll3* genotypes

We first assessed expression differences between *vgll3* genotypes in each developmental stage; two stages of *Immature1* and *2* (late spring and summer times) and *Mature* (early autumn) (Fig. 1). We found six differentially expressed genes at *Immature 1*, and 20 genes at *Immature 2* time point, as well as 25 genes at *Mature* time point between the genotypes. At *Immature 1* (late spring), all of the six differentially expressed genes identified showed higher expression in *vgll3*EE* genotype individuals (Fig. 1A). The query of molecular interactions revealed one gene, *kdm5bb*, showing direct interaction with *vgll3*, and two genes with *yap1* (*edar* and *cof2d/cfl2*) (specified with connecting lines between the genes in Fig. 1B). At *Immature 2*, we found 16 genes with higher expression in *vgll3*LL* genotype individuals and four genes showing higher expression in *vgll3*EE* genotype individuals (Fig. 1C). The predicted interactions revealed five genes (*rhoad*, *rnd3b*, *runx2*, *snaib* and *stk4*) with direct molecular interaction with *yap1* whereas one gene (*pcdh18c*) had direct interaction with *vgll3* (Fig. 1D). Furthermore, three of the differentially expressed genes (*rhoad*, *runx2* and *snaib*) formed interacting hubs (they showed the highest number of interactions of all investigated genes) and *rhoad* displayed the highest number of connections indicating its potential key functional role in this interaction network (Fig. 1D). Finally, at *Mature*, we found 21 genes with higher expression in *vgll3*LL* genotype individuals, and four genes showed higher expression in *vgll3*EE* genotype individuals (Fig. 1E). The interaction query identified five genes (*ajubab*, *lap4a/scrib*, *rnd3a*, *snai2a* and *stk3*) with direct interactions with *yap1*, whereas one gene (*arhgap6a*) had a direct interaction with *vgll3* (Fig. 1F). One of these differentially expressed genes, *snai2a*, formed an interacting hub, and also two of its interacting genes, *vdrad* and *cebpab*, appeared to make interacting hubs by connecting to three other genes (Fig. 1F).

**Figure 1:**
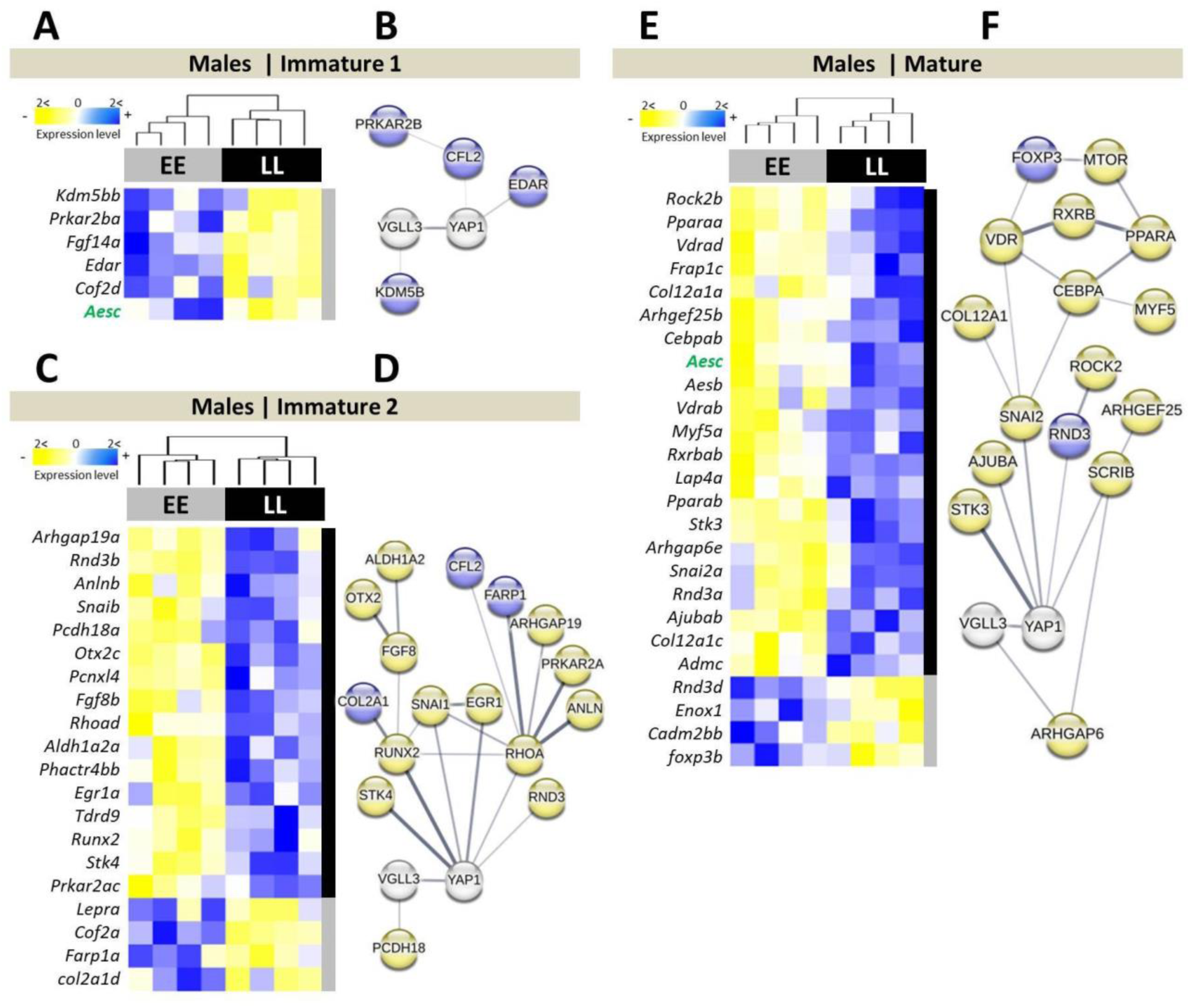
Differentially expressed genes between alternative *vgll3* genotypes and their predicted interactions in Atlantic salmon adipose tissue. Heatmaps represent differentially expressed genes between *vgll3* genotypes at three timepoints (A, C, E) and their respective predicted interactions using STRING v12 (http://string-db.org/) (B, D, F). The thickness of the connecting lines between the genes indicates the probability of the interaction. Genes colored in blue and yellow in the predicted network indicate higher and lower expression in *vgll3*EE* individuals, respectively.

### Maturation specific gene expression differences

To identify gene expression changes associated with the immature-mature transition, we compared immature individuals from both immature time points (*Immature1* and *2*) with those from the mature time point. We identified 50 DE genes between mature and immature individuals when both *vgll3* genotypes were combined, and when considering *vgll3* genotypes separately, 41 and 16 DE genes in *vgll3*LL* and *vgll3*EE* genotypes, respectively (Fig. 2A-C). Furthermore, we found a general tendency towards genes with increased expression in adipose tissue of mature males, which indicates higher transcriptional activation of the studied genes during maturation. This tendency was observed in the combined genotypes and *vgll3*LL* comparisons. In *vgll3*EE* individuals, 10 out of the 16 DE genes between immature vs mature individuals had higher expression in the matures, whereas all the DE genes in *vgll3*LL* individuals had higher expression in the matures (Fig. 2A-C). Across all three comparisons, expression patterns of three of the DE genes, *admc, col12a1a* and *pgra*, were independent of *vgll3* genotype, and all these genes showed increased expression in the mature males (Fig. 2A-D). Importantly, *yap1*, the direct opposing interactor of *vgll3*, showed a similar pattern, i.e. increased expression in mature *vgll3*LL* individuals (Fig. 2B). We further investigated potential functional/molecular interactions between DE genes showing *vgll3* genotype-specific differential expression (colored numbers in Fig. 2D). The predicted interactions between these genes revealed that among the three DE genes across all comparisons, only *pgra* had interactions with other genes in the interaction network (Fig. 2E). From the *vgll3* genotype specific comparisons, 17 and 8 DE genes within *vgll3*LL* and *vgll3*EE* individuals, respectively, were found to be connected in the interaction network (circled in blue and green in Fig. 2E). Moreover, *pgra* was connected to both the *vgll3*LL* and *vgll3*EE* interaction networks when genotypes were assessed separately through *snaia* and *foxo1c*, respectively. Other genes with a high number of interactions in the network included *ets1a*, *lats2a/b* and *wwtr1a/b* in the *vgll3*LL* interaction network, and *mob1ab*, *tead1a* and *frmd6b* in the *vgll3*EE* network (Fig. 2E).

**Figure 2:**
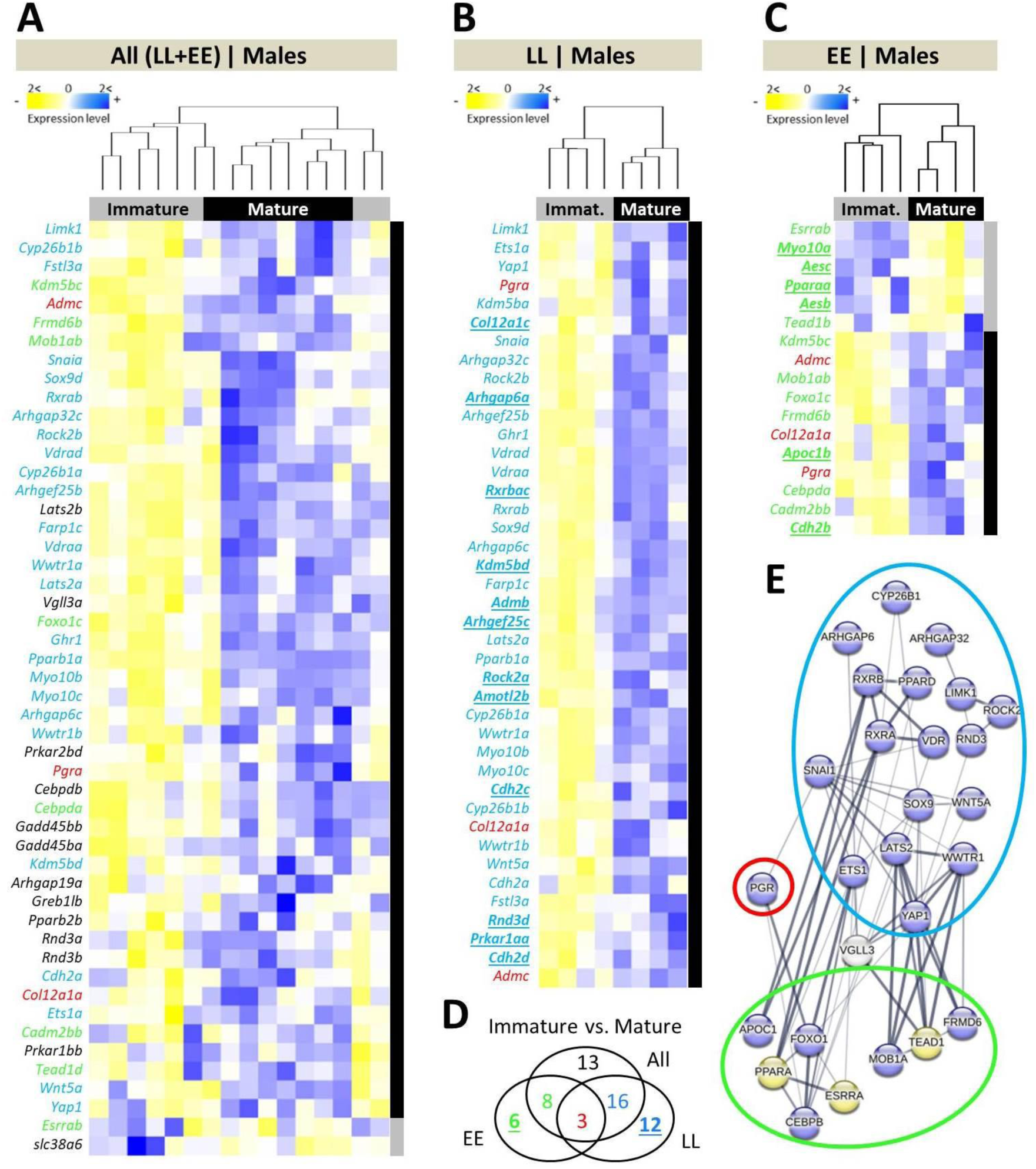
D**i**fferentially **expressed genes between immature versus mature male Atlantic salmon and their predicted interactions in adipose tissue.** Heatmaps representing differentially expressed genes between the immature versus mature males across alternative *vgll3* homozygotes (**A**), and within *vgll3*LL* (**B**) and *vgll3*EE* genotype individuals (**C**). A Venn diagram showing the numbers of differentially expressed genes overlapping between the comparisons (**D**). Predicted interactions between the overlapping genes, with green, red and blue rings indicating the genes in the Venn diagram (**E**). The thickness of the connecting lines between the genes indicates the probability of the interaction. Genes colored in blue and yellow in the predicted network indicate higher and lower expression in mature stage, respectively.

Next, we explored the gonadal development stage (using GSI) associated expression of the entire gene panel in order to identify genes with expression patterns in adipose tissue tightly linked with maturation stage. Interestingly, we found the expression of six genes; *aebp1*, *arhgap19a*, *cebpda*, *cyp26b1b*, *limk1* and *prkar2bd* to be positively correlated with GSI irrespective of *vgll3* genotype (Fig. 3A-D). Furthermore, two of these genes, *cyp26b1b* and *limk1*, displayed the most significant positive expression correlations among all the genes within samples of each genotype. In general, most of the identified correlations were positive regardless of how *vgll3* genotypes were grouped (Fig. 3A-C). Four genes amongst *vgll3*EE* genotype individuals showed negative expression correlations with increasing GSI. Among these, two genes, *esrrab* and *aesb*, were shared between *vgll3*EE* genotype and both genotypes together. Interestingly, amongst *vgll3*LL* individuals, expression of *vgll3a* was found to be among the genes positively correlated with GSI. The predicted interactions between the significantly correlated genes showed genotype-specific expression correlations (colored numbers in Fig. 3D). These revealed potential interaction networks consisting of nine and 10 genes respectively for *vgll3*LL* and *vgll3*EE* genotype individuals and four overlapping genes across all groups (Fig. 3E). The genes with the strongest and/or highest number of connections in the network include *amotl2a*, *frmd6b* and *wwtr1a* in the *vgll3*LL* interaction network, and *col1a1a* and *mob1ab* in the *vgll3*EE* network (Fig. 3E).

**Figure 3:**
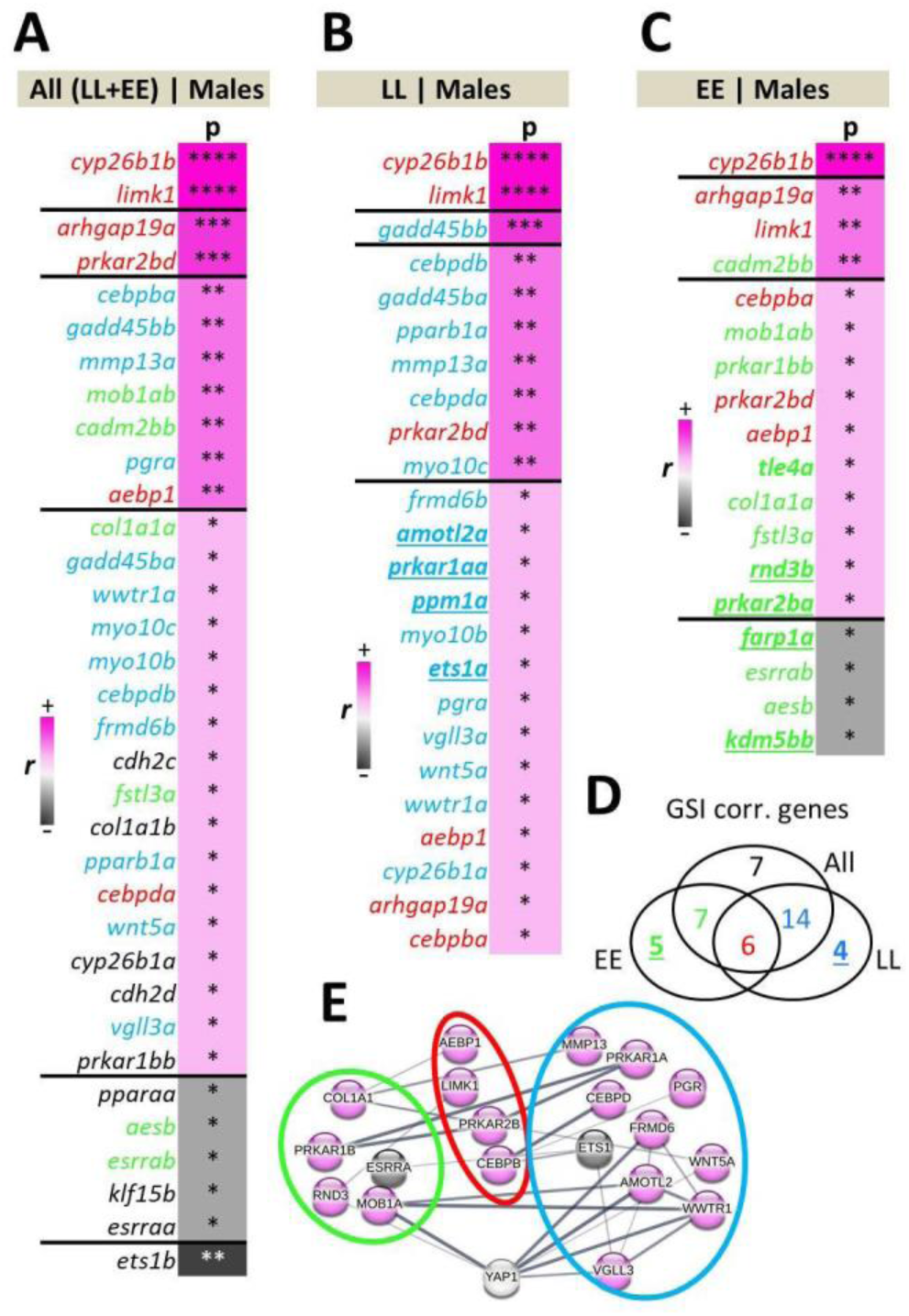
G**e**nes **showing GSI correlated expression and their predicted interactions in adipose tissue.** Ranking of significant Pearson correlations between gene expression and GSI in salmon adipose tissue across both *vgll3* genotypes (**A**), within *vgll3*LL* (**B**) and *vgll3*EE* (**C**) genotype individuals. The Venn diagram depicts the numbers of significantly correlated genes unique or overlapping between the comparisons (**D**). Predicted interactions between the overlapping genes specified with green, red and blue rings indicating genes in the Venn diagram (**E**). EE and LL indicate *vgll3*EE* and *vgll3*LL* genotypes, respectively, and p and r imply p-values (* < 0.05; ** < 0.01; *** < 0.001; **** < 0.0001) and Pearson correlation coefficient. The thickness of the connecting lines between the genes indicates the probability of the interaction.

### Identification of gene coexpression networks

In order to gain better overview of transcriptional dynamics of Hippo pathway components and their known interacting genes, we applied network-based co-expression analyses in which genotype dependent changes in each network could be tracked. To do this, we first built gene coexpression networks (GCN) in adipose tissue of each *vgll3* genotype and then investigated the preservation of the identified gene co-expression modules or GCMs (within each GCN) between the genotypes. In other words, we defined the GCN in one genotype (*vgll3*EE or vgll3*LL*) and then assessed the preservation of its modules in the other genotype (*vgll3*LL or vgll3*EE*), respectively. We identified nine GCMs for *vgll3*EE* (Fig. 4A and B) of which four, blue, turquoise, magenta and green, showed moderate preservation (Zsummary > 2) in *vgll3*LL*, i.e. most of the genes in each GCM have significant expression correlations in both genotypes. The other five GCMs (yellow, pink brown, red and black) showed a low level of preservation between *vgll3* genotypes (Zsummary < 2); (Fig. 4A).

**Figure 4.**
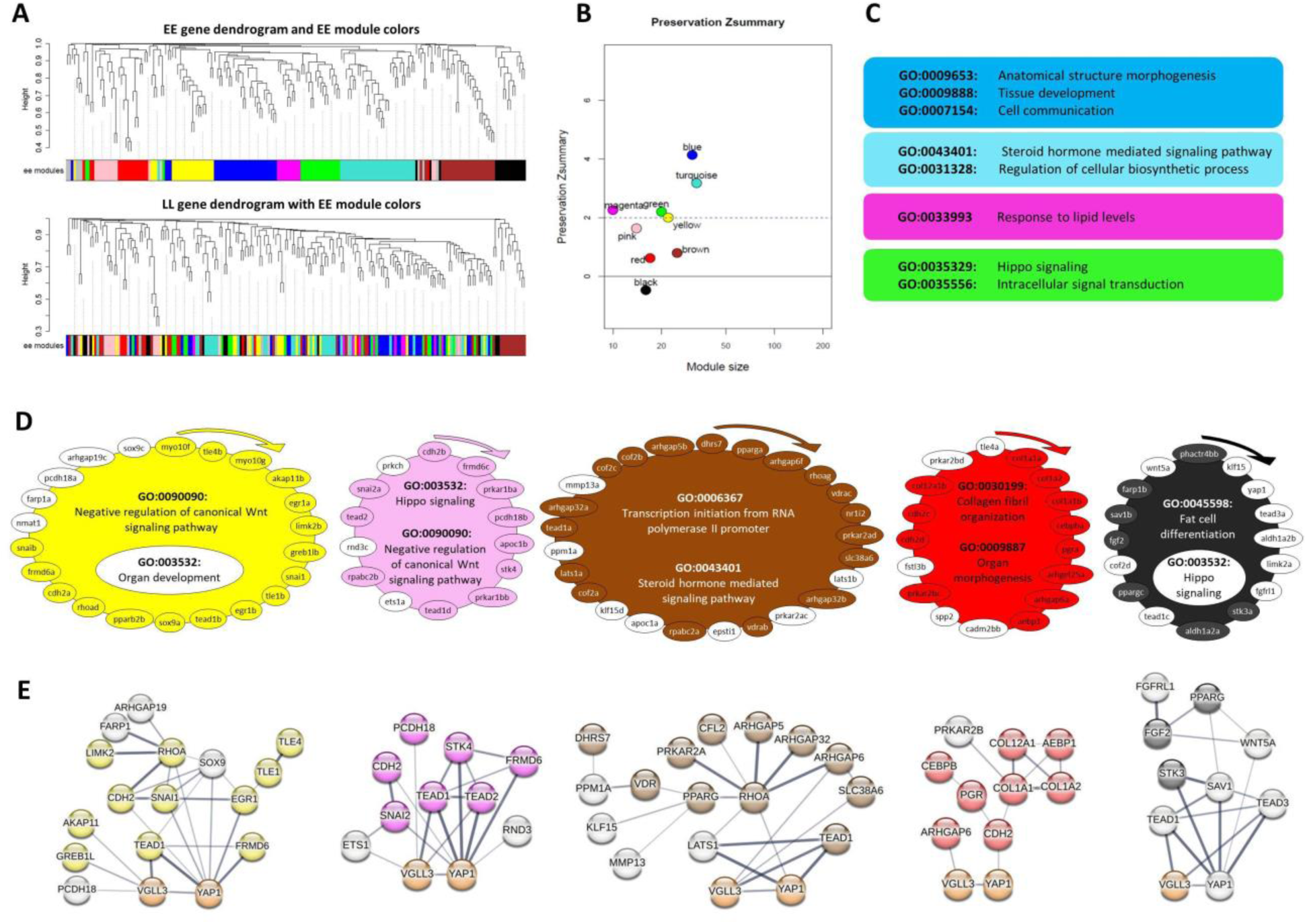
Coexpression analysis of *vgll3*EE* genotype in adipose tissue of male Atlantic salmon. (A) Visual representation of *vgll3*EE* module preservation in *vgll3*LL* individuals. The dendrograms represent average linkage clustering tree based on topological overlap distance in gene expression profiles. The lower panels of the dendrograms represent colors that correspond to the *vgll3*EE* clustered coexpression modules (GCMs). Top: *vgll3*EE* GCMs with assigned colors. Bottom: visual representation of the lack of preservation of *vgll3*EE* GCMs genes in *vgll3*LL* genotype individuals. (B) Preservation Zsummary scores in the *vgll3*LL* GCMs for *vgll3*EE* GCMs (colors represent *vgll3*EE* GCMs). Zs mmary < 2 represents lack o preservation (dotted blue line) and Zsummary between 2 and 10 implies moderate preservation. (C) The top hit GO biological processes among the most significantly enriched GOs in each of the well-preserved coexpression modules. (D) The genes in each of the identified GCMs for *vgll3*EE* genotype with least preservation in *vgll3*LL*. The genes without color in each module are those showing no preserved expression correlation in *vgll3*LL* individuals and the clockwise arrows above each GCM indicate the direction of genes with highest to lowest expression correlations with other genes within that GCM. For each of the GCMs, the top enriched GOs are represented, and GOs without color were no longer enriched after removal of the genes without colors. (E) Predicted interactions between the genes within each of the less preserved GCMs. Different levels of thickness in the connecting lines between the genes indicates the probability of the interaction.

GO Biological Process enrichment analysis was conducted for each of the GCMs separately in order to identify the key biological processes genes in each GCM are involved in. Enriched GO terms for the moderately preserved GCMs included tissue development and morphogenesis, steroid hormone signaling and Hippo signaling pathways as well as response to lipid levels (Fig. 4C). In the low preserved GCMs, we found coupling of Hippo pathway with fat cell differentiation in the black GCM and with negative regulation of canonical Wnt pathway in the pink GCM (Fig. 4C). The black GCM showed the least preservation between the genotypes with most of its genes showing no coexpression preservation in *vgll3*LL* genotype individuals (genes lacking color in each GCM in Fig. 4D). Removal of unpreserved genes in the yellow and black GCNs led to loss of significance of one GO in each GCM; organ development in the yellow and Hippo signaling in the black GCM (non-coloured GOs in Fig. 4D). The removal of the unpreserved genes in the other GCNs did not change their enriched GOs (Fig. 4D).

Knowledge-based interactome prediction analysis using genes within each GCM was applied in order to identify potential interactions between the genes as well as hub genes with highest number of interactions. The prediction of interactions between the genes within each GCM revealed that except in the red GCM, all the other GCMs had at least one gene among those not showing coexpression preservation had direct interaction with *vgll3/yap1* (represented with lines directly connecting the non-colored genes with *vgll3/yap1*; Fig. 4E). This was more pronounced in the black GCM where four genes (*sav1b*, *tead1c*, *tead3a* and *wnt5a*) had direct interactions with *vgll3/yap1* (Fig. 4E). Importantly, *yap1* itself lost coexpression preservation in the *vgll3*LL* GCM, indicating the involvement of yap1, the major inhibitor of the Hippo pathway, in differences between the *vgll3* genotypes in the lowest preserved GCM.

In *vgll3*LL* individuals, we found seven GCMs and among them only the blue GCM showed a low level of preservation (Zsummary < 2) compared to *vgll3*EE* individuals (Fig. 5A and B). The turquoise GCM was the largest with 74 co-expressed genes and also with the highest preservation level between the genotypes (Zsummary > 7). Interestingly, the turquoise GCM included genes encoding components of the Hippo signaling pathway and genes that negatively regulate the canonical Wnt pathway (Fig. 5C), indicating potential interactions between Hippo and Wnt pathway in *vgll3*LL* individuals. We observed such connections between these pathways in a GCM of *vgll3*EE* as well (the pink GCM in Fig. 4D), however, the number of genes in the pink GCM (14 genes) was much less than the turquoise GCM (76 genes) (Fig. 4B and Fig. 5B). This may indicate more extensive connections between regulators of Wnt pathway and components of Hippo pathway in *vgll3*LL* individuals. On the other hand, the blue GCM, which was the least preserved GCM in *vgll3*LL*, appeared as the second largest with 37 genes (Fig. 5B and D). This may indicate that the Hippo pathway is more affected and/or plays a more extensive role in the adipose tissue of *vgll3*LL* individuals. The blue GCM displayed coupling of genes within Hippo signaling pathway with organ development and morphogenesis (Fig. 5D), but removal of unpreserved genes led to no significant enrichment of the GO linked to tissue morphogenesis (non-colored genes and GO in Fig. 5D). The prediction of interactions between the genes within the blue GCM revealed that three genes that lost their coexpression preservation, *mob1aa*, *stk3b* and *tead2*, had direct interactions with *vgll3/yap1* (Fig. 5E).

**Figure 5.**
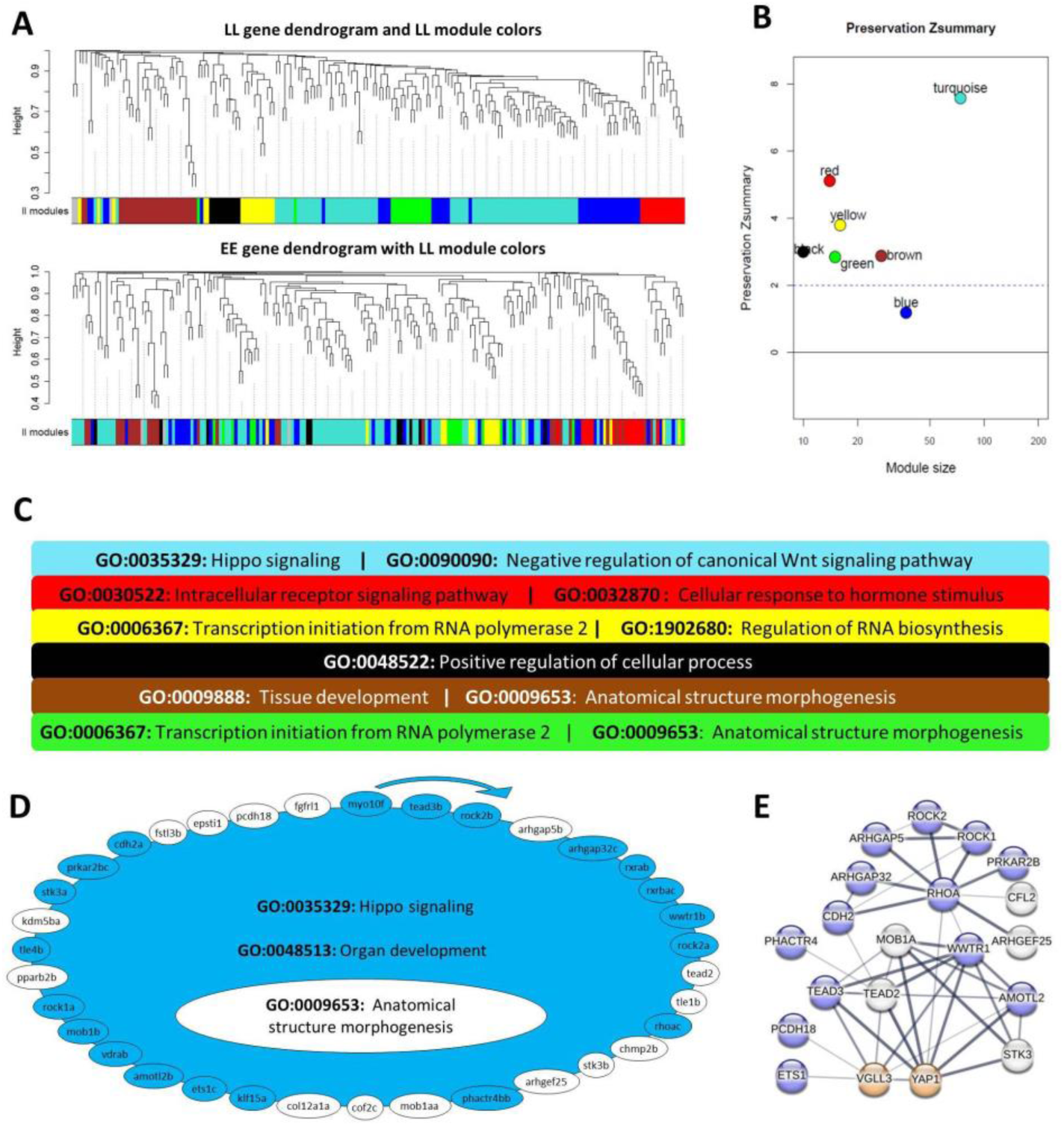
Coexpression analysis of *vgll3*LL* genotype in adipose tissue of male Atlantic salmon. (A) Visual representation of *vgll3*LL* module preservation in *vgll3*EE* individuals. The dendrograms represent average linkage clustering tree based on topological overlap distance in gene expression profile. The lower panels of the dendrograms represent colors that correspond to the clustered coexpression modules (GCMs). Top: *vgll3*LL* GCMs with assigned colors. Bottom: visual representation of preservation of *vgll3*LL* module genes in *vgll3*EE* genotype. (B) The colors represent identified *vgll3*LL* GCMs. Preservation of genes found in *vgll3*LL* GCMs in the *vgll3*EE* GCMs was calculated and Zs mmary < 2 represents lack o preservation (dotted blue line). Zsummary between 2 and 10 implies moderate preservation. (C) The most significant gene ontologies (GO) enriched in each of the well-preserved coexpression modules. (D) The genes in each of the identified coexpression modules for *vgll3*LL* genotype with least preservation in *vgll3*EE*. The genes without color in each module have lost their preserved expression correlation in *vgll3*EE* individuals and the clockwise arrows above each GCM indicate the direction of genes with highest to lowest expression correlations with other genes within that GCM. For each of the module, the most significant enriched GOs are represented, and GOs without color have lost their significance after removal of the genes without colors. (E) Predicted interactions between the genes within each of the less preserved GCMs. The connecting lines between the genes with different levels of thickness indicate the probability of the interaction.

## Discussion

Understanding the molecular basis of life-history traits and their interaction with the environment is crucial to predict organismal fitness in future climates. In this study, we aimed to uncover transcriptional effects of alternative maturation age genotypes of the *vgll3* gene, a major determinant of age at sexual maturity in Atlantic salmon (Barson et al., 2015) and also a main co-factor of the Hippo pathway (Hori et al., 2020), in adipose tissue of this species. To do this, we investigated the expression differences of genes encoding adipogenesis related factors, Hippo pathway components and their associated interacting partners/signals between *early* (E) or *late* (L) maturing *vgll3* alleles using a custom-made Nanostring platform. This allowed us to predict the regulatory dynamics of Hippo pathway signaling in the adipose tissue of Atlantic salmon that has a seasonal breeding strategy tightly associated with lipid metabolic status. Thus, the study of distinct *vgll3* genotypes provides understanding of the molecular links among lipid metabolism, Hippo pathway activity, and sexual maturation – a life-history trait of high ecological importance. Our findings not only imply that *vgll3* and its associated pathway (Hippo) have extensive effects on transcriptional changes in adipose tissue in relation to sexual maturation of Atlantic salmon, but also suggest *vgll3*’s role in linking adipogenesis and seasonal, likely maturation related, changes in this species.

### Gene expression patterns at the immature stage reflect higher lipid storage capacity of *vgll3*EE* individuals in the spring

A recent study showed a link between *vgll3* genotypes and lipid storage in the liver of immature males of Atlantic salmon during seasonal changes (from spring to autumn) (House, Debes, Holopainen, et al., 2023). A possible hepatic scenario was proposed based on the differences observed between *vgll3* genotypes; i.e., *vgll3*EE* individuals storing larger lipid droplets in earlier/warmer season (spring), whereas *vgll3*LL* individuals storing in a later/colder season (autumn) in their liver (House, Debes, Holopainen, et al., 2023). Our adipose tissue gene expression results support this proposed hepatic scenario in the immature stage. However, the *vgll3* genotypic associations were reversed in the adipose of mature stage individuals. Our results suggest that *vgll3*LL* individuals are unlikely to store larger lipid droplets during the spring due to increased expression of a RhoA encoding gene (*rhoad*) in their adipocytes. RhoA is a well-known key suppressor of adipogenesis and lipid droplet storage in mammalian cells (Meyers, Zayzafoon, Douglas, & McDonald, 2005), and has also been suggested to be inhibited by *vgll3* dependent activation of Hippo pathway (Kurko et al., 2020). Vgll3 itself has been reported as an inhibitor of adipocyte differentiation in mice (Halperin et al., 2013) and linked with body condition variation in Atlantic salmon (Debes et al., 2021). Thus, our finding here of higher expression of the adipogenesis suppressor, *rhoad*, in *vgll3*LL* individuals is consistent with the hepatic scenario suggested by House et al., 2023, indicating potentially similar regulatory pattern in the liver and adipose. Furthermore, we found *vgll3*LL* individuals to have higher expression of a paralogue gene (*rnd3b*) encoding RND3, a key lipolysis factor (Dankel et al., 2019). Lipolysis is a process in which lipid droplets are utilized (the opposite to lipogenesis) (Ducharme & Bickel, 2008), and thus this finding may also supports the hepatic scenario proposed by (House et al., 2023), whereby immature *vgll3*EE* individuals store larger lipid droplets during the spring.

### Gene expression patterns at the mature stage reflect higher lipid storage capacity of *vgll3*LL* individuals in the early autumn

In mature males, the higher expression of major adipogenic factors *snai2a*, *cebpab* and *ajubab,* as well as reduced expression of lipolysis factors; *rnd3a* and *pparaa* in *vgll3*LL* individuals indicates an opposite pattern in mature compared to immature males (i.e., lower lipid storage capacity of *vgll3*LL* individuals in immature individuals). The observed pattern in mature individuals suggests that *vgll3*LL* individuals store larger lipid droplets in autumn compared to *vgll3*EE* individuals, suggesting an opposite shift between the genotypes from immature to mature status. A key component of the interacting hub among the differentially expressed genes between the genotypes in the mature stage was a paralogue of *Snai2*/*Slug* gene (*snai2a*) (Fig. 1E and F). Interestingly, *Snai2* is known as an inducer of adipocyte differentiation and lipid accumulation (Pérez-Mancera et al., 2007). Similarly, a paralo e o /EBPα encodin ene (*cebpab*) with predicted interaction with Snai2 displayed higher expression in *vgll3*LL* individuals (Fig. 1F). /EBPα is a major ind cer o late sta e adipocyte differentiation, and in mammalian cells, its expression is directly promoted by the transcriptional co-regulator Ajuba (Yan et al., 2022). Consistently, a paralogue gene encoding Ajuba (*ajubab*) was also among the genes with higher expression in *vgll3*LL* individuals (Fig. 1F). In mammals, *Ajuba* is a direct transcriptional target gene for Yap1 during cell proliferation and differentiation, and increased activity of Yap1 induces *Ajuba* expression (Lange et al., 2015). Moreover, in contrast to the immature stage, we found lower expression of *rnd3a* in *vgll3*LL* compared to *vgll3*EE* individuals at the mature time point (Fig. 1F). RND3 is an inhibitor of Rho kinase (ROCK) signaling and an important regulator of lipid metabolism in adipocytes via induction of lipolysis (Dankel et al., 2019). In agreement with the reduced expression of the lipolysis factor, *rnd3a*, we also observed reduced expression of *pparaa* in *vgll3*LL* individuals which encodes a major receptor activatin lipolysis (PPARα) in adipose and liver (Guzmán et al., 2004). Combined, these expression patterns suggest higher lipid storage capacity of *vgll3*LL* individuals in the early autumn at mature stage.

### Gene expression patterns reflect reduced lipid storage capacity at the mature stage in both *vgll3* genotypes but with less Hippo pathway dependency in *vgll3*EE*

Comparison of expression patterns between immature and mature stages suggest reduced adipogenesis and increased lipolysis in the mature males, and subsequently, the utilization of lipid droplets to invest energy in sexual maturation. Comparing all immature and mature individuals, only one gene encoding progesterone receptor (*pgra*) and a paralogue gene of its downstream target, adrenomedullin (*admc*) were differentially expressed independently of *vgll3* genotype across all the immature versus mature comparisons. In mammals, it has been recently demonstrated that PGR can act as inhibitor of adipogenesis (X. Liu et al., 2021) and also be a stimulator of lipolysis (Zhang et al., 2021). We observed similar patterns between mature and immature individuals within each *vgll3* genotype based on the expression pattern of genes involved in lipogenesis and lipolysis. However, it seems that the reduced adipogenesis (and lipogenesis) of mature individuals is the result of different mechanisms within each genotype. In *vgll3*LL* individuals, the vast majority of DE genes exhibited an increase in expression at the mature stage (including *lats2* and *yap1*). This may suggest that lipolysis in mature *vgll3*LL* individuals might be mediated through increased expression of a major Hippo pathway kinase, *lats2*, which is also known to promote lipolysis (El-Merahbi et al., 2020). In *vgll3*EE* mature individuals, other regulators independent of the Hippo pathway such as FOXO1 and ESRRα (encoded by *foxo1c* and *esrrab*) appeared to be the mediators of the potential processes of increased lipolysis and reduced lipogenesis. FOXO1 has a critical role in transcriptional induction of the major adipocyte lipolysis enzyme (adipose triglyceride lipase ATGL) (Chakrabarti & Kandror, 2009). Estrogen-related receptor α (ESRRα) is a n clear receptor that promotes adipocyte differentiation by regulating the expression of various adipogenesis-related genes (Ijichi et al., 2007). Hence, the higher expression of *foxo1c* and the lower expression of *esrrab* in *vgll3*EE* mature compared to immature individuals could be another indicator of reduced adipogenesis and increased lipolysis in the mature males. However, a gene encoding C/EBP-β (*cebpba*), an early stage inducer of pre-adipocyte proliferation was also upregulated in the mature *vgll3*EE* individuals (Merrett, Bo, Psaltis, & Proud, 2020). Taken together, this suggests that the adipose tissue of the mature males in this genotype might have higher number of undifferentiated pre-adipocytes, however, histological studies are required to approve this hypothesis. The perpetuation of undifferentiated stage in pre-adipocytes might be also linked to the inhibitory role of *vgll3* during adipogenesis, which mainly blocks the terminal stage of adipocyte differentiation (Halperin et al., 2013).

### Gene expression-GSI correlations suggests reduced lipid storage capacity at the mature stage in both *vgll3* genotypes

To gain more insight into the molecular details of the effects of maturation on adipose gene expression, we also conducted correlation analysis between gene expression and gonadal maturation level (GSI score). A notable gene amongst those with the strongest positive correlations with gonadal maturation level across genotypes was a gene encoding LIM domain kinase 1 (*limk1*), which is an inhibitor of adipocyte differentiation through modulation of intracellular structural organization required during the differentiation (L. Chen, Hu, Qiu, Shi, & Kassem, 2018). Interestingly, *limk1* is a direct downstream target of PGR in humans (Mazur et al., 2015), thus this result is consistent with the potential role of PGR in mature individuals generally showing lower lipidogenesis and higher lipolysis capacity (see above). This suggests reduced adipogenesis in the mature males of both genotypes. However, clear expression pattern differences between the *vgll3* genotypes were observed as well. For instance, in mature *vgll3*LL* individuals, we found a positive correlation between *vgll3a* expression and gonadal maturation suggesting a more direct role of *vgll3a* itself in inhibition of adipogenesis during maturation in this genotype (Halperin et al., 2013). In *vgll3*EE* individuals, we observed negative correlation between *esrrab* expression and gonadal maturation suggesting diminished ESRRα-mediated adipocyte differentiation, as ESRRα is a known ind cer o adipocyte differentiation (Ijichi et al., 2007). Hence, this may indicate differential expression o ESRRα to be an important mechanism for reduced adipogenesis in *vgll3*EE* individuals (Fig. 2), and also suggests that the reduced adipogenesis in this genotype is less dependent on the Hippo pathway. A summary of all abovementioned findings in respect to salmon maturation, seasonal effects, and adipogenesis is depicted in Fig 6.

**Figure 6.**
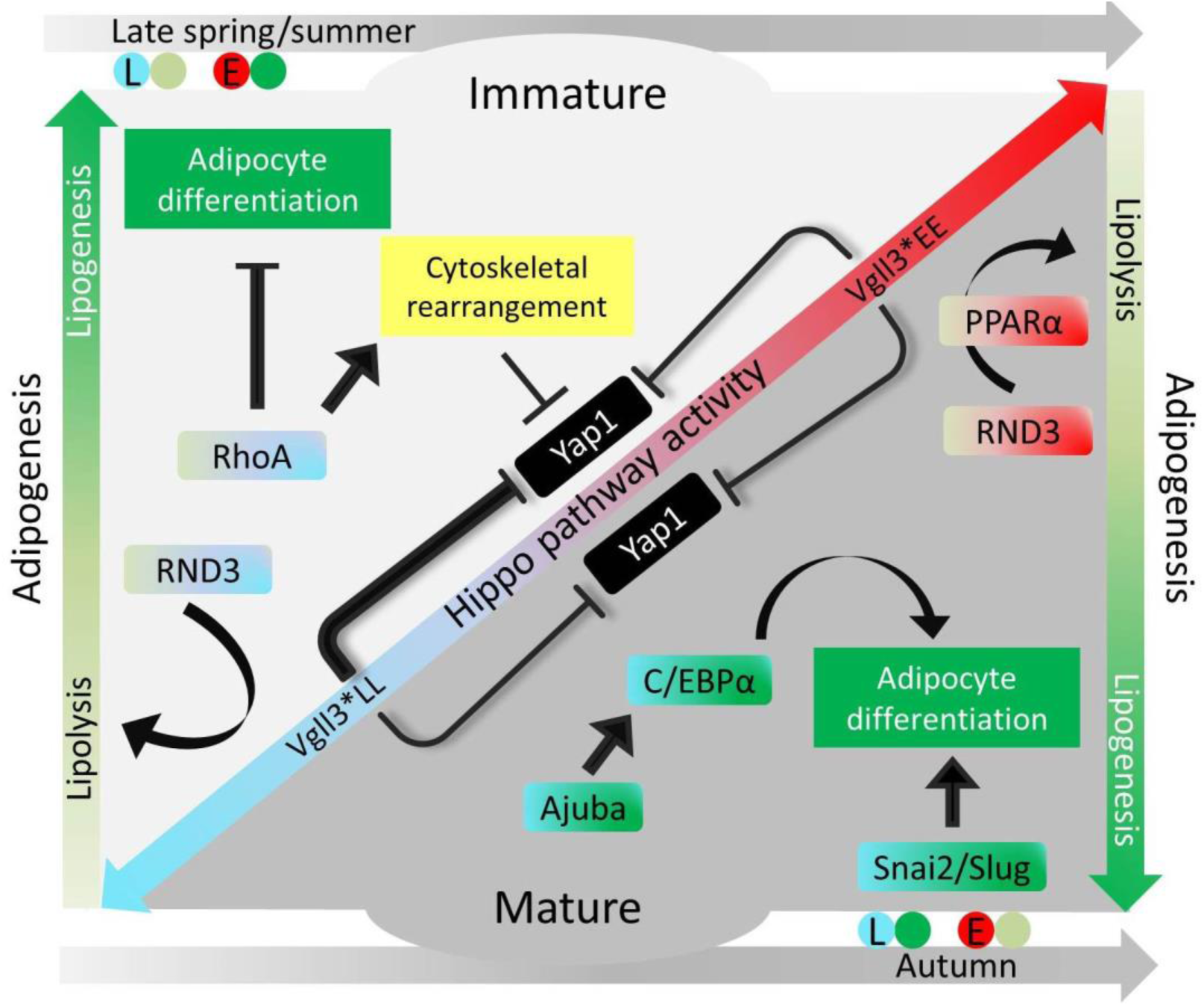
A schematic representation of the main findings linking vgll3 genotype and Hippo pathway components to different processes in the adipose tissue of Atlantic salmon. Colors indicate Hippo pathway modes of activity through the late and early maturation *vgll3* genotypes (*vgll3*LL*: light blue and *vgll3*EE*: red), as well as distinct adipogenic/lipogenic (dark green) and lipolytic (light green) processes. Genes are color coded according to *vgll3* genotype (blue or red) and lipogenesis/lysis statuses (dark or light green) indicating higher expression in the respective genotype and/or the resulting lipogenesis/lysis outcome of their higher expression. Arrow heads indicate activation whereas blocked head implies inhibition.

### Gene expression network analysis suggests differing Hippo pathway associations in *vgll3*EE* and *vgll3*LL* individuals

Hippo pathway is emerging as a key molecular signal in balancing adipocyte proliferation versus differentiation, and the activity of Yap has been found to be indispensable during adipogenesis (Ardestani et al., 2018). We conducted gene co-expression network analysis to identify genotype-specific differences within the network of co-expressed genes in adipose tissue, and found that in *vgll3*LL* individuals, there was a lack of association between expression of the Hippo pathway components and genes involved in adipocyte differentiation. In the GCM identified in *vgll3*EE*, with the lowest preservation level between the genotypes (the black GCM in Fig. 5), we observed a lack of a link between the GO terms related to fat cell differentiation from the GO of the Hippo pathway. Interestingly, several genes encoding major components of Hippo pathway were among the genes that lacked coexpression preservation in *vgll3*LL* (*tead1c*, *tead3a*, *sav1b* and *yap1*). Sav1, an inhibitor of Yap, has been shown to play a main role in enhancing adipocyte differentiation in mammals (Ardestani et al., 2018). Thus, these findings in the black GCM suggest that there are distinct transcriptional patterns of Hippo pathway components between the *vgll3* genotypes in relation to adipogenesis, which might be a result of different *sav1b* and *yap1* regulatory functions in these genotypes. However, a direct regulatory connection between Sav1 and Vgll3 has not been investigated in any species. In the yellow GCM, we found no association between the GO term related to negative regulation of Wnt signaling and the GO of organ development. Strikingly, among the genes in this GCM showing lack of expression preservation between the genotypes was a negative regulator of Wnt signaling, *cdh2* (encoding N-cadherin), which is known to enhance adipogenesis by inhibiting Wnt signal (Haÿ et al., 2014). Moreover, *cdh2* is a well-known regulator of organ development (García-Castro, Vielmetter, & Bronner-Fraser, 2000), and also a downstream transcriptional target of vgll3 (together with *pcdh18a* which also lacks coexpression preservation in the yellow GCM) (Simon, Thézé, Fédou, Thiébaud, & Faucheux, 2017). Finally, *cdh2* appeared to be critical for structural/mechanical regulation of Hippo pathway during adipose tissue development (Nardone et al., 2017).

In *vgll3*EE* individuals, we found a lack of association between expression of the Hippo pathway components and genes involved in structural morphogenesis. Among the identified GCMs in *vgll3*LL* genotype individuals, only one GCM containing genes involved in Hippo pathway, organ development and structural morphogenesis (depicted in blue in Fig. 5) showed a low level of preservation between the genotypes. The adipose tissue morphogenesis and structural remodeling are major contributors to metabolic, functional and adaptive changes in this tissue (Choe, Huh, Hwang, Kim, & Kim, 2016; F. Liu, He, Wang, Zhu, & Bi, 2020). When the blue GCM was compared to its structure in *vgll3*EE* genotype individuals, there was a lack of co-expression preservation for genes involved in tissue morphogenesis and remodeling such as *cof2c* (*CFL2*), *col12a1a* and *fstl3b* (H. jian Chen et al., 2023; Galkin et al., 2011; Li et al., 2023) and adipogenesis such as *fgfrl1* (Wang et al., 2020). The preservation loss in this GCM resulted in the dissociation of the GO term related to structural morphogenesis from the other GOs (Hippo pathway and organ development) (Fig. 5). In addition, there was also an absence of coexpression preservation in *vgll3*EE* individuals for several members of Hippo pathway (*mob1aa*, *tead2a* and *stk3b*). Since the pathway itself is critical for adipose tissue remodeling and morphological modification (Lecoutre et al., 2022; Shen et al., 2022), the lack of correlations with genes involved in morphogenesis is most likely a downstream consequence of transcriptional differences between the genotypes in these three key components of the Hippo pathway. Reciprocally, morphological/structural changes at the intra-and extra cellular levels can mediate their effects on adipose tissue by regulating Hippo pathway signaling (L. Liu et al., 2022; Seo & Kim, 2018). These findings suggest that the differing role of the Hippo pathway between *vgll3* genotypes potentially results in distinct structural/morphological changes in the adipose tissue of the two *vgll3* genotypes in salmon, and these differences are likely to be dependent on Yap/Taz transcriptional activity.

## Conclusions

This study provides important molecular evidence linking ecological change (seasonality), physiological process (lipid metabolism/storage), and genotype-linked life-history strategy variation (sexual maturation). Our findings provide gene expression evidence supporting the previously proposed scenario in which Atlantic salmon immature males exhibiting genotype specific lipid profiles with *vgll3*EE* individuals increasing lipid storage between spring and autumn while *vgll3*LL* individuals do the opposite. Furthermore, such a pattern seems to be reversed (i.e., *vgll3*EE* individuals decreasing lipid storage) in the mature individuals during autumn. At the molecular level, our results suggest that certain components of the Hippo signaling pathway (e.g., *yap1*, *sav1*, *lats2*, etc.) are potentially major drivers of the differing effects of *vgll3* genotypes on the adipose tissue gene expression. However, more detailed functional assessments are required to better validate these suggested differences.

## Acknowledgements

We thank N. Piavchenko, S. Andrew, O. Andersson, T. Aykanat, Y. Czorlich, J. Erkinaro, A. House, M. Lindqvist, N. Lorenzen, O. Mehtälä, K. Mobley, J. Moustakas-Verho, O. Ovaskainen, S. Papakostas, N. Parre, A. Ruokolainen, V. Pritchard, K. Salminen, M. Sinclair-Waters, S. Tillanen, and K. Zueva for help related to gamete stripping, sample processing, tagging, genotyping, phenotyping or fish husbandry, and the staff at the Natural Resources Institute Finland (Luke) hatchery in Taivalkoski hatchery for help during spawning.

## Author Contributions

EPA, CRP, JPV, JK and PVD conceived the study; JPV, PVD and CRP reared and sampled the fish; CRP and JE provided resources; AR, EPA and JPV performed experiments; EPA, JPV, PS and JK developed methodology and analyzed the data; EPA, JPV and CRP interpreted results of the experiments; EPA, JPV and CRP drafted the manuscript, with EPA having the main contribution, and all authors approved the final version of manuscript.

## Ethical approval

Animal experimentation followed European Union Directive 2010/63/EU under license ESAVI/35841/2020 granted by the Animal Experiment Board in Finland (ELLA).

## Data availability

All the gene expression data generated during this study are included in this article as supplementary file.

Supplementary Data: Expression data and statistical analysis.

